# XAI-MRI: An Ensemble Dual-Modality Approach for 3D Brain Tumor Segmentation Using Magnetic Resonance Imaging

**DOI:** 10.1101/2024.11.19.624437

**Authors:** Ahmeed Suliman Farhan, Muhammad Khalid, Umar Manzoor

## Abstract

Brain tumor segmentation from Magnetic Resonance Images (MRI) presents significant challenges due to the complex nature of brain tumor tissues. This complexity makes distinguishing tumor tissues from healthy tissues difficult, mainly when radiologists perform manual segmentation. Reliable and accurate segmentation is crucial for effective tumor grading and treatment planning. In this paper, we proposed a novel ensemble dual-modality approach for 3D brain tumor segmentation using MRI. Initially, individual U-Net models are trained and evaluated on single MRI modalities (T1, T2, T1ce, and FLAIR) to establish each modality’s performance. Subsequently, we trained U-net models using combinations of the best-performing modalities to exploit the complementary information and improve segmentation accuracy. Finally, we suggested the ensemble dual-modality by combining the best-performing two pre-trained dual-modalities models to enhance segmentation performance. Experimental results show that the proposed model enhanced the segmentation result and achieved a Dice Coefficient of 97.73% and a Mean IoU of 60.08% on the BraTS2020 dataset. The results illustrate that the ensemble dual-modality approach outperforms single-modality and dual-modality models. This study shows that ensemble dual-modality models can help improve the accuracy and dependability of brain tumor segmentation based on MRI. Our code publicly available at: https://github.com/Ahmeed-Suliman-Farhan/Ensemble-Dual-Modality-Approach

## INTRODUCTION

Brain tumors represent one of the most severe and complex challenges in the medical field (Muhammad et al., 2021). They arise from abnormal growth of cells within the brain or inside the skull (Biratu et al., 2021). It is one of the most dangerous diseases and one of the leading causes of death in various countries.(Alqhtani et al., 2024). The most common type of brain tumors is gliomas, which are considered a challenge for both doctors and researchers because of diversity and difficulty in diagnosing and treating them. It is estimated that about 80,000 people in the United States are diagnosed with brain tumors each year, and the majority of them have gliomas (Zhou, 2023; Thakkar et al., 2020). Gliomas are classified into two main groups according to their grade: high-grade gliomas (HGG) and low-grade gliomas (LGG). Determining the grade has a significant role in planning the treatment. (Sun et al., 2023). Although low-grade gliomas are less aggressive than high-grade gliomas, they can develop into higher-grade gliomas if there is no treatment in time, which increases the severity of the disease and complicates treatment (Bogdańska et al., 2017; Claus et al., 2015). Therefore, early diagnosis of brain tumors is essential and increases the chances of patients survival and treatment (Al-Zoghby et al., 2023; Saeedi et al., 2023). Diagnosis of brain tumors begins with MRI because it is the most efficient tool for imaging brain tissue. In addition, MRI provides a three-dimensional view of the brain, which helps doctors better diagnose the tumor(Abd-Ellah et al., 2024).

Magnetic resonance imaging (MRI) is a form of multimodality imaging. MRI generates images of different contrasts because protons in the tissues vary in their relaxation rates (Zhan et al., 2022; Zhou et al., 2019). Different modalities of MRI images, such as T1-weighted, T1-weighted images with contrast enhancement (T1c), FLAIR, and T2-weighted help doctors better view the tumor (Tandel et al., 2023). For example, T2 and FLAIR modalities target on the edema area surrounding the tumor while T1 and T1c focus on the tumor area (Zhou et al., 2019). Figure 1 shows the different modalities of brain MRI images. These different modalities help doctors accurately analyze and diagnose the tumor and develop treatment plans (Hammad et al., 2023; Sailunaz et al., 2023). However, brain tumors from MRI images can be difficult to recognise and segment due to high variability in tumour shape, size and location (Almufareh et al., 2024). In addition, manual segmentation is time-consuming and prone to errors (due to variability in interpretations) and requires an expert radiologist. Therefore, there is an urgent need to develop an automatic system for segmenting brain tumors from MRI images. This system can help segment and diagnose tumors accurately and efficiently (Karim et al., 2024; Hussain and Shouno, 2024).

**Figure 1.**
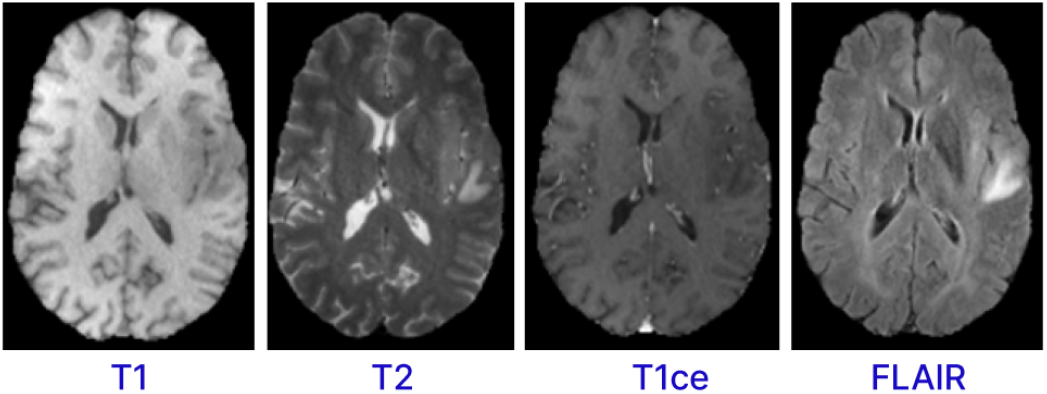
MRI Sequences for Brain

Accurately segmenting brain tumours from magnetic resonance imaging (MRI) data is important to diagnosis. However, segmentation is complicated due to the different brain tumors types and the difficulty distinguishing between infected and healthy cells, highlighting the urgent need for an automated segmentation system. In recent years, deep learning, especially the U-Net architecture, has shown promising results in medical image segmentation (Zhang et al., 2024a). Because each modality provides limited information, single-modality MRI methods often do not perform well in brain tumor segmentation. Although multiple MRI modalities can enhance segmentation accuracy, most current segmentation models do not integrate features from different modalities. This study suggests a new ensemble dualmodality method based on the U-Net model. This method aims to improve segmentation performance by training and integrating bimodal U-Net models, making the method for brain tumor segmentation more accurate.The study shows that the proposed ensemble dual-modality model performed better than the single-modality and multi-modality models. The main contributions of this study are as follows:

- An ensemble dual-modality module is proposed to combine two pre-trained dual-modality models. These pre-trained models use dual modalities to improve segmentation accuracy.
- This study evaluates the results of single-input and multi-input input segmentation by training and testing the U-net model on both input types.
- The study proposed a novel approach for preprocessing 3D MRI images and cropping the region of interest. Accurately identifying and cropping the region of interest reduces the complexity of the segmentation and ensures that the segmentation model focuses on the most relevant areas of the brain.
- The proposed method was rigorously evaluated using the BraTS2020 dataset. The study shows the effectiveness of the ensemble dual-modality model in accurately segmenting brain tumors.

## LITERATURE REVIEW

In recent years, researchers have become focused on medical image analysis, and specifically brain tumour segmentation from MRI images. Various deep learning-based brain tumour segmentation models have been proposed. Deep learning methods have attention due to their ability to learn features automatically. U-net architecture has emerged as a popular choice in medical image segmentation among deep learning methods due to its ability to extract features efficiently and segment tumors. Several studies have explored the use of U-net model for brain tumor segmentation. For example, (Zhou, 2024) suggested a novel technique for multi-modal brain tumor segmentation using U-net. The proposed method uses disentangled representation learning and region-aware contrastive learning. Disentangled representation learning separates the fused feature into separate factors corresponding to various tumour parts. Contrast learning facilitates the extraction of feature representations relevant to tumor regions. The proposed method is tested on BraTS 2018 and BraTS 2019. The proposed approach demonstrated better performance than the other current methods.

(Fang and Wang, 2022) proposed a dual-path network to segment brain tumors more effectively. The dual-path network can also take advantage of the data in different modalities to help improve brain tumor segmentation from MRI images. The proposed model was trained and run on the BraTS 2015 dataset, and its performance is efficient.

(Zhou, 2023) proposed a novel multimodal model for brain tumour segmentation from MRI images. This approach could slice brain tumors when one or more modalities are missing and can retrieve the missing modalities. The proposed model utilizes features from multiple modalities for brain tumour segmentation. Latent feature learning also extracts multimodal latent correlations. The proposed approach was tested and evaluated on the BraTS 2018 dataset and achieved promising segmentation results when one or more modalities were missing.

(Montaha et al., 2023) They suggested a 2D U-Net-based approach to segmenting brain tumors from 3D MRI data. The proposed method was trained and tested on the BraTS2020 dataset, and the results obtained 93.1% DSC and 99.41% accuracy.

(Yousef et al., 2023) proposed a novel Deep Learning-based Brain Tumor Segmentation architecture called Bridged-U-Net-ASPP-EVO for Brain tumor segmentation from MRI scans. It includes spatial hierarchical pooling (ASPP) and an advanced normalization layer. The proposed approach was tested and evaluated using the BraTS 2020 and BraTS 2021 data. The test results on the BraTS 2020 dataset showed an average of 0.78, 0.8159 and 0.9073 for ET, TC and WT, respectively, and HD95% of 21.684, 15.941 and 5.37. The test results on the BraTS 2021 dataset had averaged DSC of 0.8434, 0.8594 and 0.9187 for ET, TC and WT, respectively, and average HD95% of 11.669, 141887 and 5.3687.

(Feng et al., 2024) suggested a new way to represent frequencies to reduce feature loss in the segmentation model, primarily when brain tumor detection is encoded and decoded. This method, called MLU-Net, integrates frequency representation techniques and Multilayer Perceptron (MLP)-based techniques into a lightweight U-Net architecture. MLU-Net does a better job of medical image segmentation tasks while keeping the computer’s processing power high by using frequency representation and MLP-based methods. In experiments on brain tumor segmentation, the proposed approach demonstrates significant efficiency enhancements, achieving reductions in parameters and computational workload to merely 1/39 and 1/61 of those required by the U-Net model. In addition, the proposed approach outperforms U-Net by enhancing both Dice and Intersection over Union (IoU) metrics by 3.37% and 3.30%, respectively.

(Hammer Håversen et al., 2024) proposed a novel approach for 3D segmentation using a selfsupervised and self-querying framework integrated into the U-Net architecture. Unlike traditional methods that rely solely on labeled data for supervision, QT-UNet leverages self-supervision to learn from unlabeled data, reducing the need for annotated samples. The experimental results when using the brain tumor segmentation (BraTS 2021) dataset demonstrate the effectiveness of the proposed approach, where it obtained a Hausdorff Distance of 4.85 mm and an average Dice score of 88.61. (Zhu et al., 2024) proposed a method that leverages multi-modality MRI data to enhance spatial information and correct boundary shapes for more accurate tumor segmentation. Their approach comprises three modules: border shape correction (BSC), spatial information enhancement (SIE), and modality information extraction (MIE). The three modules comprise an end-to-end 3D brain tumor segmentation model, acting on deep convolutional networks’ input, backbone, and loss functions (DCNN). The experiments demonstrated promising results, which were validated on BraTS2017, BraTS2018, and BraTS2019 datasets.

(Ranjbarzadeh et al., 2024) proposed a brain tumor segmentation framework using four modalities of MRI image types. The proposed model is based on convolutional neural networks (CNNs) and Improved Chimp Optimization Algorithm (IChOA). Initially, all four MRI modality types (T1, T2, T1ce, Flair) are normalized to identify potential tumor regions. Then, IChOA is used to select features using a Support Vector Machine (SVM) classifier. Finally, these features are fed into the proposed CNN model for tumor segmentation. Using IChOA contributes to feature selection and hyperparameter optimization of the proposed CNN model. Experimental results on the BRATS 2018 dataset show that the proposed model achieved a Precision of 97.41%, recall of 95.78%, and dice score of 97.04%.

(Zhang et al., 2024b) proposed a novel approach named ETUNet (Efficient Transformer Enhanced UNet) for 3D brain tumor segmentation. This method integrates transformer modules into the UNet architecture to leverage their efficiency in capturing long-range dependencies and enhancing feature representations. By incorporating transformers, ETUNet aims to improve the segmentation accuracy and efficiency compared to traditional UNet models. Through experimental evaluations on BraTS-2018 and BraTS-2020 datasets, ETUNet obtained average Dice Similarity Coefficient (DSC) scores of 0.854 and 0.862 and 95th percentile of the Hausdorff Distance (HD95) values of 6.688 and 5.455 on two datasets, respectively.

While the studies mentioned have showcased the effectiveness of U-net for brain tumor segmentation, they often face limitations in leveraging the complementary information from multiple MRI modalities with a multi-path model. Additionally, existing approaches may suffer from robustness and reduced generalization capability. To address these limitations, our proposed ensemble Dual-Modality approach integrates different MRI modalities and ensemble Dual paths to enhance segmentation accuracy. By leveraging the strengths of each modality and combining them through an ensemble framework, we aim to improve the robustness and efficiency of brain tumor segmentation while mitigating the challenges associated with single-modality approaches.

## PROPOSED METHODOLOGY

The methodology proposed in this research is an ensemble dual-modality approach for brain tumor segmenting from MRI data. Figure 2 shows all stages of the proposed technology. In the first stage, the BraTS2020 dataset was chosen to evaluate the effectiveness of the proposed segmentation method. The second stage of pre-processing the dataset includes identifying and cropping the region of interest and resizing it, then partitioning it into testing, training, and validation sets. After that, the third stage is feature extraction and segmentation of the brain tumor, comprising three key steps: Step one trains and evaluates the U-net model on each MRI modality separately, including T1, T2, T1ce, and FLAIR. The second step involves training and evaluating the U-net model using multi-modalities. The third step combined two pre-train models with input modalities T2+T1ce and T1ce+FLAIR, which obtained the best results from the previous step in creating an ensemble dual-modality segmentation model. Together, these stages form a comprehensive framework for enhancing the efficiency of brain tumor segmentation. The following sections explain each stage in more detail, including its impact on brain segmentation performance.

**Figure 2.**
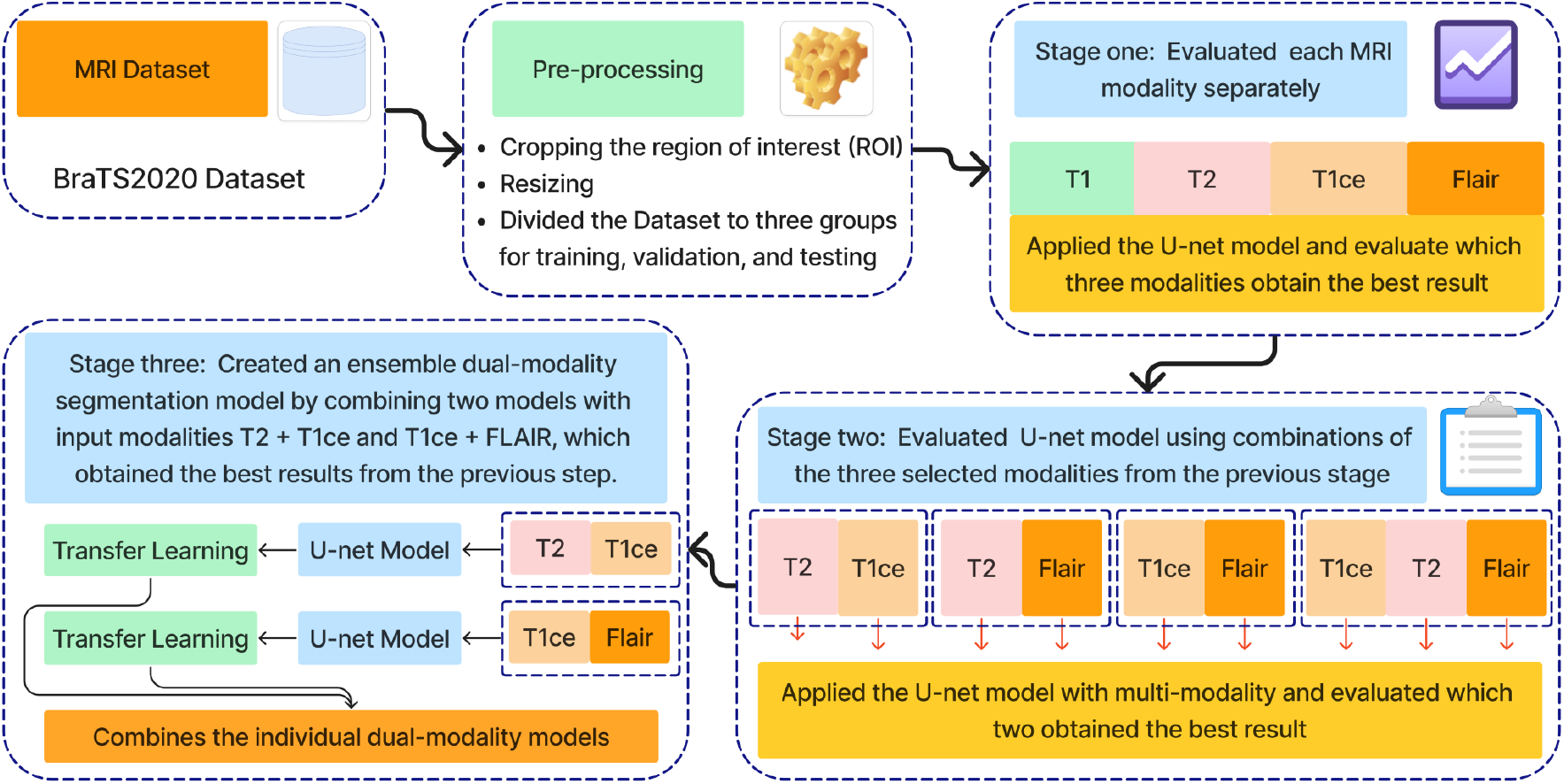
Workflow of the proposed approach.

**Figure 3.**
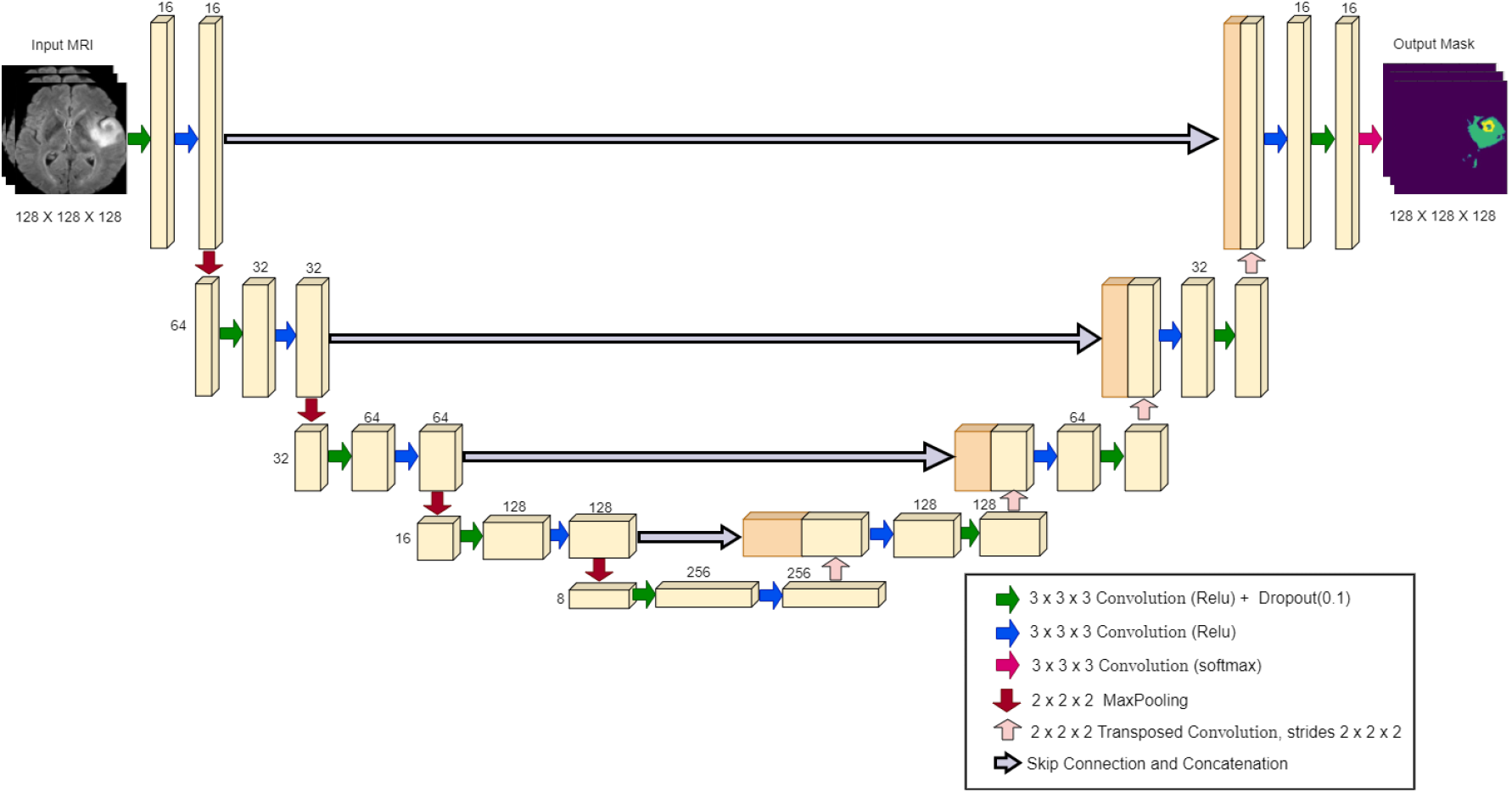
U-Net Model Architecture.

### U-Net Model Architecture

The U-Net architecture is a convolutional neural network widely used in biomedical image segmentation. It was introduced by (Ronneberger et al., 2015), and is characterized by a symmetrical “U” shape, consisting of a contracting path (encoder) and a dilating path (decoder). The encoder captures the context through convolutional layers and max-pooling layers, gradually reducing the spatial dimensions as depth increases. The decoder then upsamples the features, using the transferred convolutions, and connects them to the corresponding encoder features through skip connections (Ibtehaz and Rahman, 2020; Montaha et al., 2023). Figure 5 illustrates the specific configuration of the 3D U-Net architecture utilized in our study, highlighting its layers and connections.

### Data sets

This study used the Brain Tumor Segmentation 2020 (BraTS2020) dataset to train and test the proposed approach. This dataset contains 494 3D image subjects, divided into 369 subjects for training and 125 for validation. Each subject includes MRI scan data across multiple modalities: T1-weighted, T1 post-contrast (T1ce), T2-weighted, and FLAIR (fluid-attenuated inversion recovery). Each modality has a resolution of 240 × 240 in the axial plane, paired with a depth of 155 slices. The training dataset also contains ground truth labels that have been manually reviewed by neuroradiologists. Therefore, in this study, we used these 369 subjects for training and testing the model. The annotations include (tumor core (TC), whole tumor (WT), enhanced tumor (ET)) (Menze et al., 2014; Bakas et al., 2017, 2018)

### Pre-processing

The preprocessing of the BraTS2020 dataset includes cropping the region of interest (ROI). The ROI cropping step is important and improves the performance of segmentation models by making the model focus on the relevant parts of the brain. This study used 369 subjects from the BraTS2020 dataset to train and test the model. These 369 subjects include the ground truth labels.

To define and crop the ROI for each subject, we chose the T1 modality of MRI images because this modality demonstrates the contrast between the brain and the surrounding edges. Slice number 77 was chosen from the T1 modality, which includes a depth of 155 slices. As the middle slice, this typically offers a central view of the brain and the tumour, ensuring that the cropped region in all other slices encompasses the whole brain. This slice was used to calculate the coordinates x1, x2, y1, and y2, which are critical to determining the ROI. Subsequently, we reduced the depth from 155 to 128 slices by selecting slices from 13 to 141. The next step is cropping all slices by applying the same calculated coordinates value on all 128 slices of the MRI across all modalities: T1, T2, T1ce, FLAIR, and the mask. After that, the cropped images were resized to 128×128 pixels, resulting in a standardized input size of (128, 128, 128) for the segmentation model. Figure 4 shows these steps in detail.

**Figure 4.**
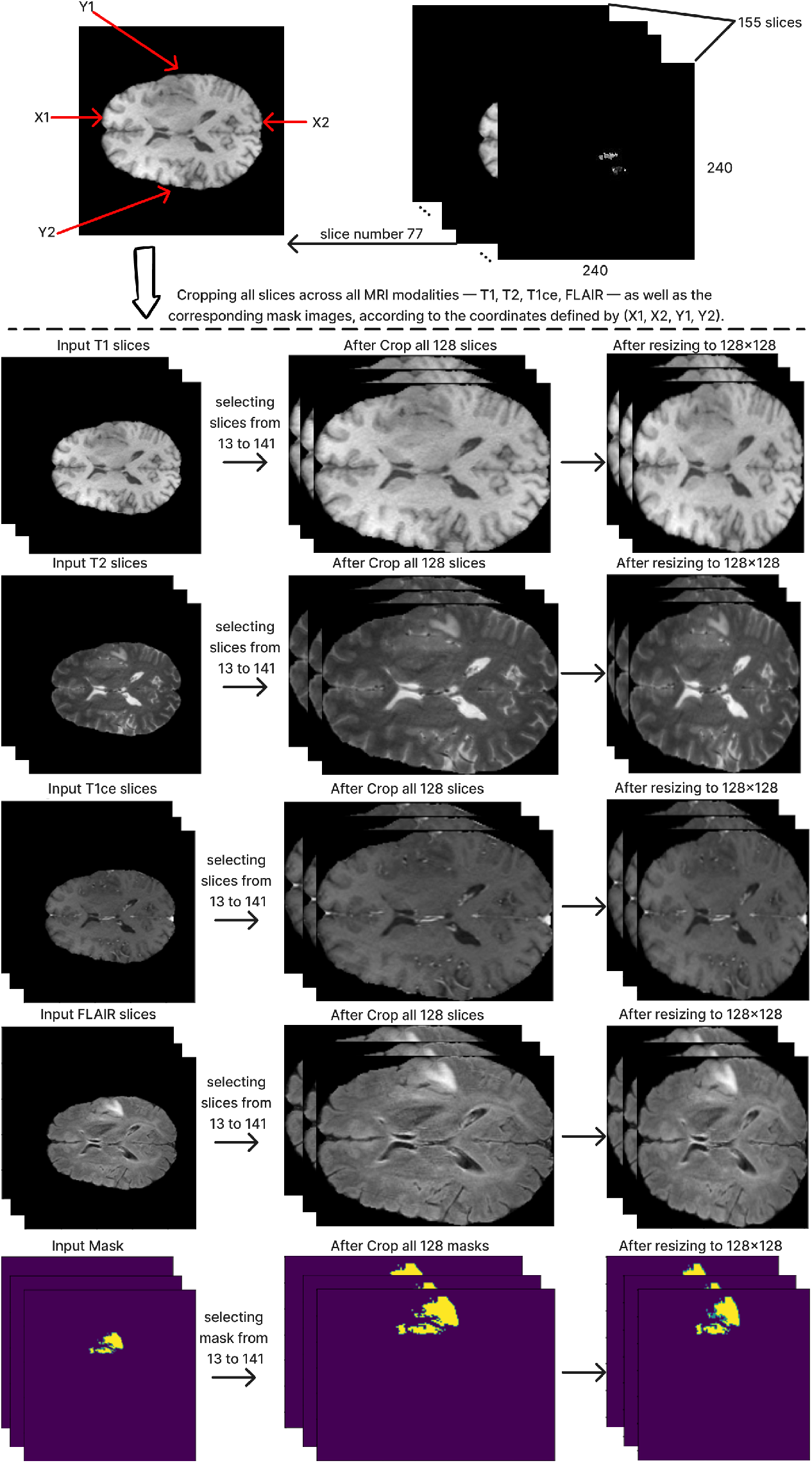
Description Preprocessing steps

Finally, the dataset was divided into three groups for training, validation, and testing. The distribution was 70% of the dataset for training, 15% for validation, and 15% for testing.

### Ensemble Dual-Modality Method

This paper introduces an ensemble dual-modality approach to enhance brain tumor segmentation capabilities and help clinical decision-making. Individual Unet models are trained and evaluated on each MRI modality separately, including T1, T2, T1ce, and FLAIR. This step determines each method’s essential tumor segmentation performance. as shown in Figure 5. The T2 and T1ce Flair methods were selected, which obtained the best results for the next step.

**Figure 5.**
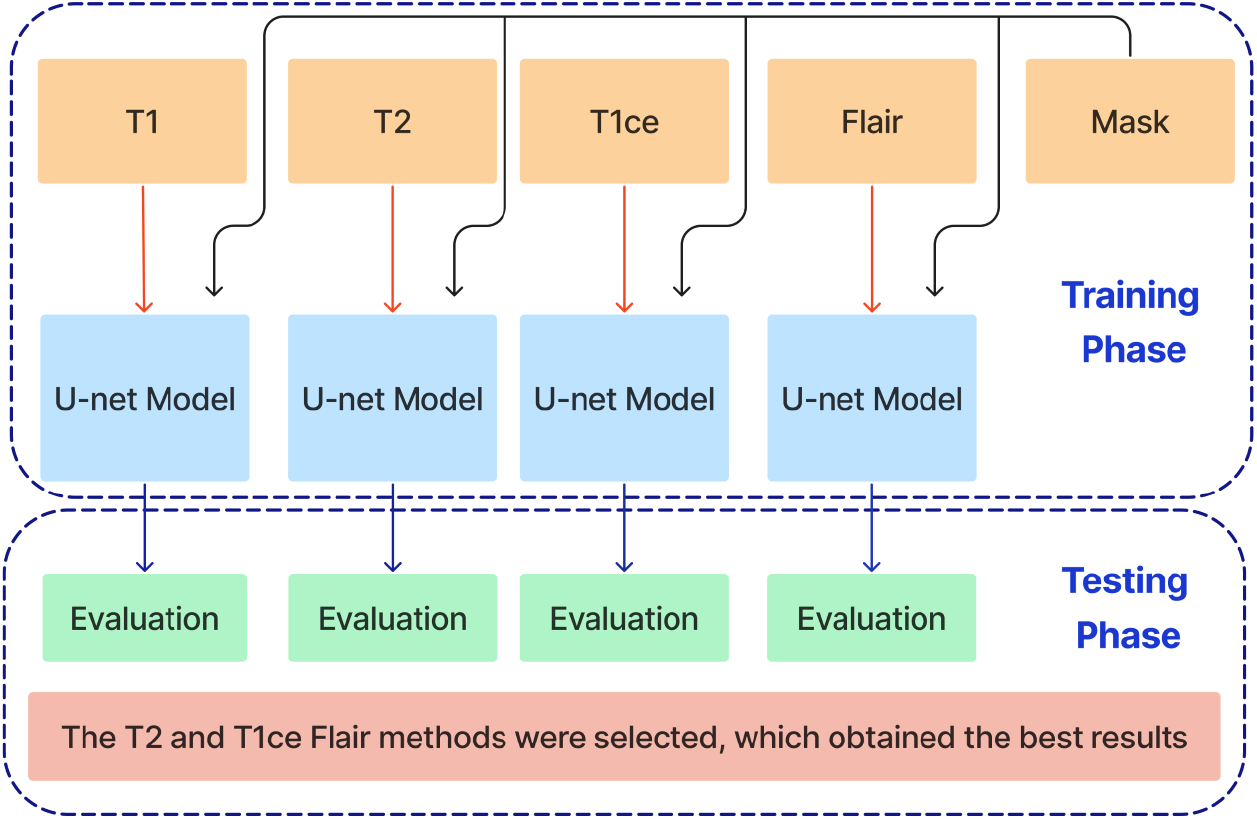
Training and Evaluation of individual Unet models on separate MRI modalities (T1, T2, T1ce, and FLAIR) for brain tumor segmentation.

In the next stage, Unet models are trained using combinations of the three selected modalities, which obtained the best results from the previous step. This method trains models on multi-input modalities, such as T2 + T1ce, T2 + FLAIR, T1ce + FLAIR, T2 + T1ce + Flair, as shown in Figure 6. This method exploits complementary information in different modalities to enhance segmentation accuracy. Using multiple modalities simultaneously can yield more robust segmentation results because each MRI modality captures unique aspects of tumor characteristics.

**Figure 6.**
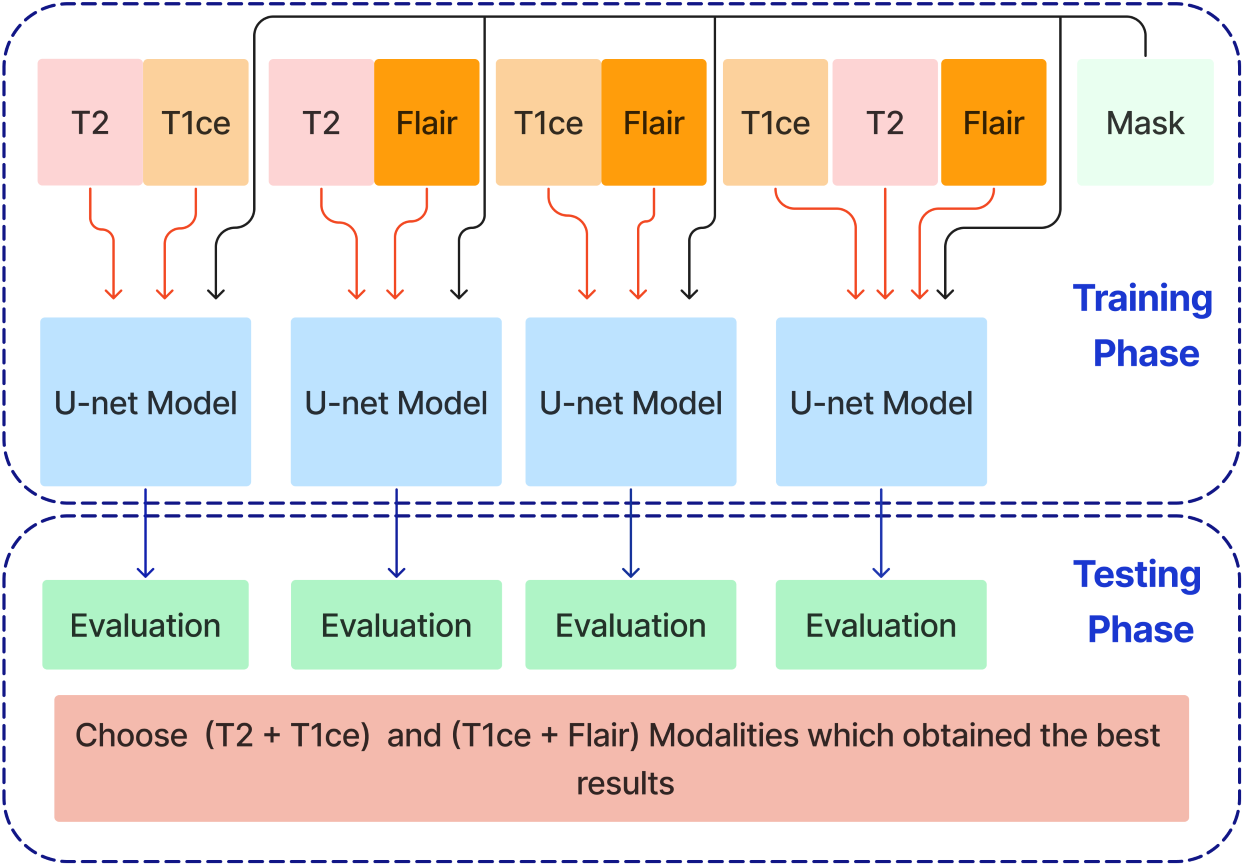
Training and Evaluation Unet models using combinations of selected MRI modalities (T2 + T1ce, T2 + FLAIR, T1ce + FLAIR, T2 + T1ce + FLAIR) for enhanced brain tumor segmentation.

After training the dual modality models, the pre-trained models are combined. We chose the two models with input modalities T2 + T1ce and T1ce + FLAIR that obtained the best results from the previous steps to create an ensemble dual-modality segmentation model. This ensemble process combines the individual dual-modality models by removing the output layer from each model and adding a Concatenate Layer to concatenate their feature maps. They were followed by an additional convolutional layer with (a Relu activation function, a filter size of (3 × 3 × 3), a stride size of 1 and an output shape of 16) Afterwards, an output layer with a softmax activation function as shown in Figure 7. The Ensemble Dual-Modality method aims to harness the complementary strengths of each modality, leading to a more comprehensive and accurate segmentation of brain tumors. The Ensemble Dual-Modality approach integrates multiple MRI modalities and ensemble the Unet deep learning model to improve the accuracy and reliability of brain tumor segmentation.

**Figure 7.**
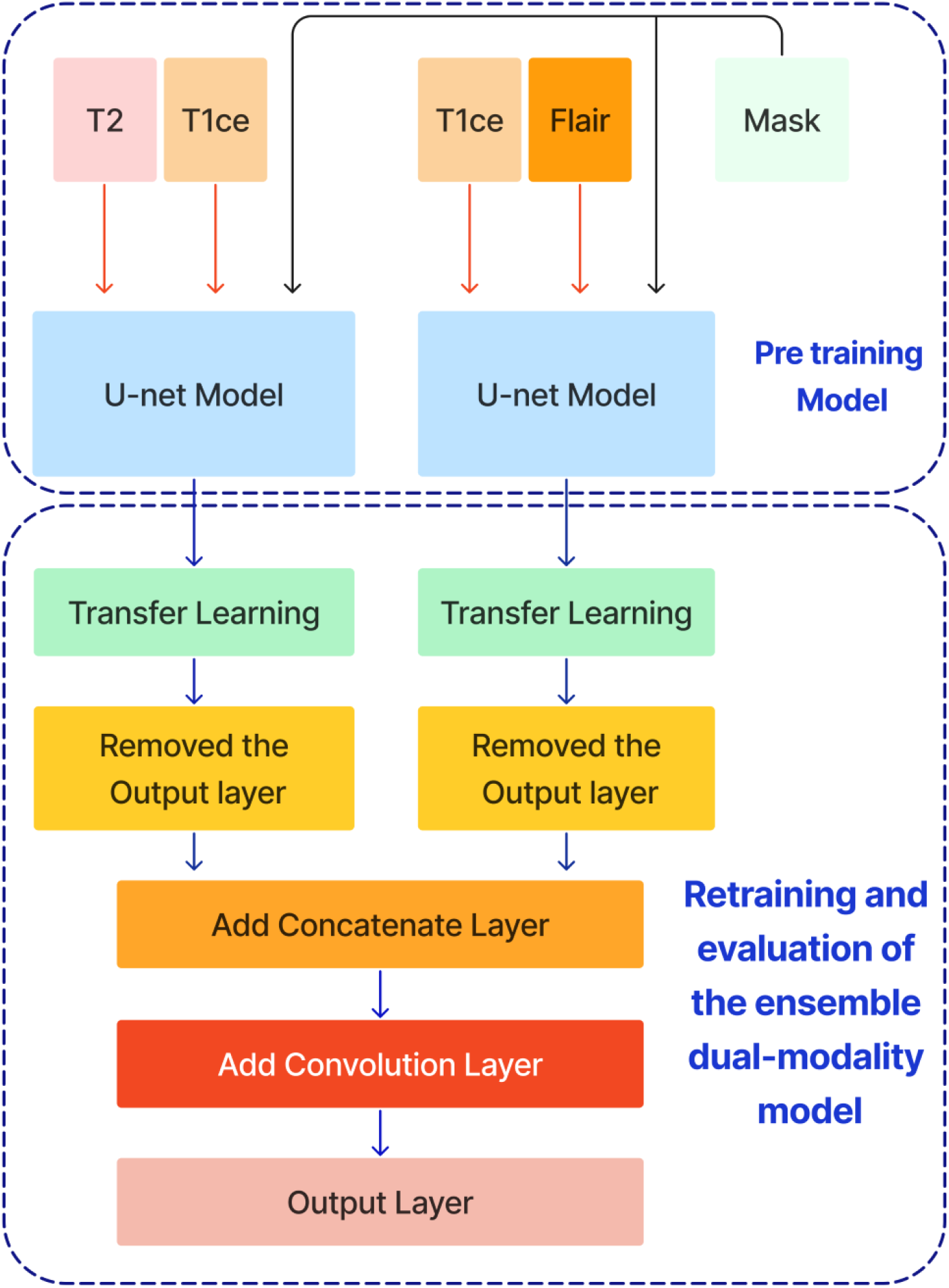
The Ensemble Dual-Modality Segmentation Model by combining the best-performing dual-modality models (T2 + T1ce and T1ce + FLAIR).

## EXPERIMENT RESULTS

This section comprehensively overviews the specific hyperparameters used in the experiments. It also describes the performance metrics used to demonstrate the effectiveness of the ensemble dual-modality model in segmenting brain tumors from MRI images. It then details the ensemble dual-modality testing results on the BraTS 2020 dataset.

### Experimental Setup

The proposed model was implemented using Python 3.8.0, Keras 2.6.0, and TensorFlow 2.6.0 libraries. The experiments were performed on a device with the following specifications: OS (Linux: Ubuntu LTS (GNU/Linux 5.15.0-97generic x86 64)), Processor (Intel(R) Xeon(R) CPU E52680 v4 @ 2.40GHz), RAM (128 GB) and GPU (05:00.0 VGA compatible controller: Matrox Electronics Systems Ltd. MGA G200e [Pilot] ServerEngines (SEP1) (rev 05))

### Hyperparameters

Before starting the training process, basic hyperparameters must be set, including batch size, number of epochs, and learning rate. For this task, the batch size was 4, the number of epochs was 40, and the learning rate was 0.0001. There was also an early stopping point for stopping training if validation loss did not improve after five consecutive epochs. This strategy helps to avoid overfitting and improves computational efficiency by aborting training when further development is not likely. These weights with the smallest validation loss were saved and used to test the approach.

### Performance metrics

To evaluate the performance of our brain tumor segmentation models, we used several key metrics: accuracy, mean intersection over union (Mean IoU), Dice similarity coefficient (DSC), precision, sensitivity, and specificity. These measures show you how the proposed approach performs from different aspects (Verma et al., 2024; Jyothi and Singh, 2023; Renugadevi et al., 2023).

#### 1. Accuracy

The segmentation accuracy is based on the proportion of correctly classified pixels (tumor and non-tumor) in comparison with total pixels. Accuracy, although helpful for generalising performance, it can be misleading in imbalanced data, such as segmenting brain tumors, where pixels from non-tumor often trump tumour pixels.

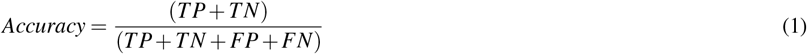

#### 2. Mean Intersection over Union (Mean IoU)

The mean IoU is a very important metric for evaluating the accuracy of the segmentation. It measures the mean overlap between the predicted and ground truth mask. The equation gives the mean IoU:

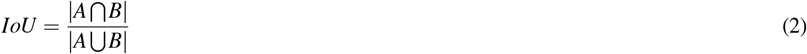

#### 3. Dice Similarity Coefficient (DSC)

The DSC is another important metric that measures the similarity between the predicted segmentation and the ground truth. It is defined as twice the overlap area divided by the total number of pixels in the predicted and ground truth masks. The equation gives DSC:

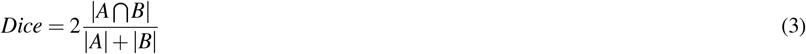

where *A* is the set of pixels in the predicted segmentation, and *B* is the set of pixels in the ground truth.

#### 4. Precision

Precision measures the accuracy of the model’s positive predictions, defined as the ratio of actual positive pixels to the sum of true positive and false positive pixels. It indicates the number of predicted tumor pixels that belong to the tumor.

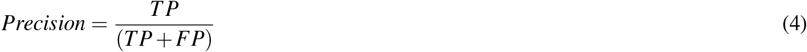

#### 5. Sensitivity (Recall)

Sensitivity or recall measures a model’s ability to correctly identify all true positive pixels. It is the ratio of true positive pixels to the sum of true positive and false negative pixels.

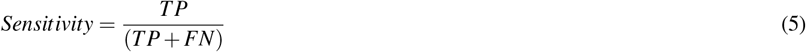

#### 6. Specificity

Specificity measures a model’s ability to correctly identify all true negative pixels. The ratio of true negative pixels to the sum of true negative and false positive pixels.

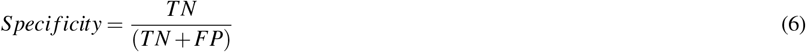

While all of these metrics are important, we primarily focused on the mean IoU and Dice Similarity Coefficient (DSC) in our study. These metrics are more useful for evaluating segmentation tasks because they directly measure the overlap between predicted and actual tumor regions, providing a clearer picture of the model’s performance in accurately defining tumor boundaries.

## Results

In this section, we trained and tested the proposed model using the BraTS2020 dataset, which includes T1, T1c, T2, and FLAIR modalities. This experiment was divided into three scenarios: single modality, multi-modality, and ensemble dual-modality. Table 1 and Figure 8 show the summary of the test results for all scenarios.

**Table 1.**
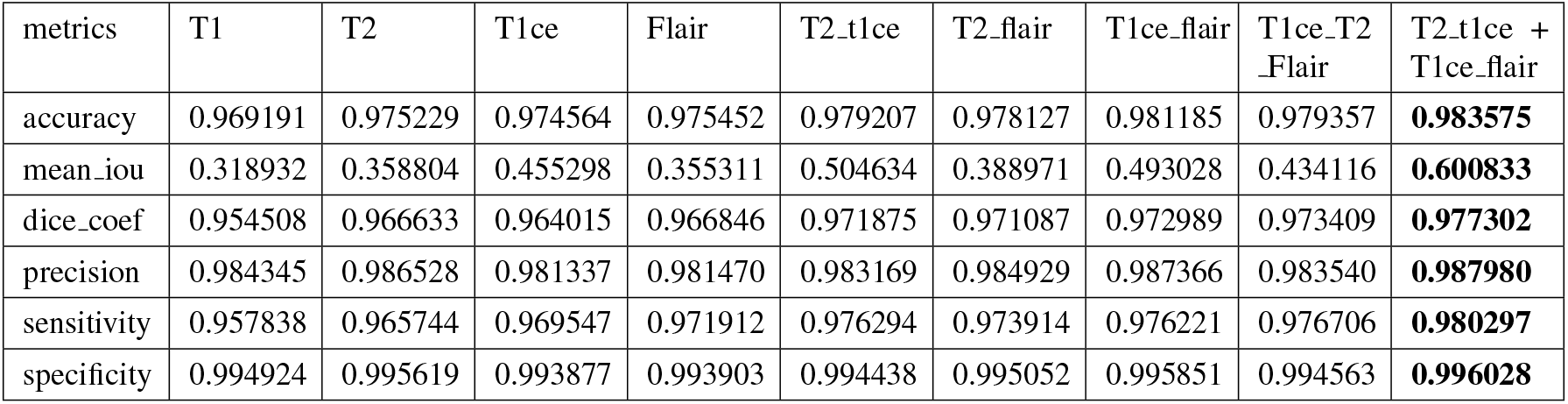
Results for all scenarios.

**Figure 8.**
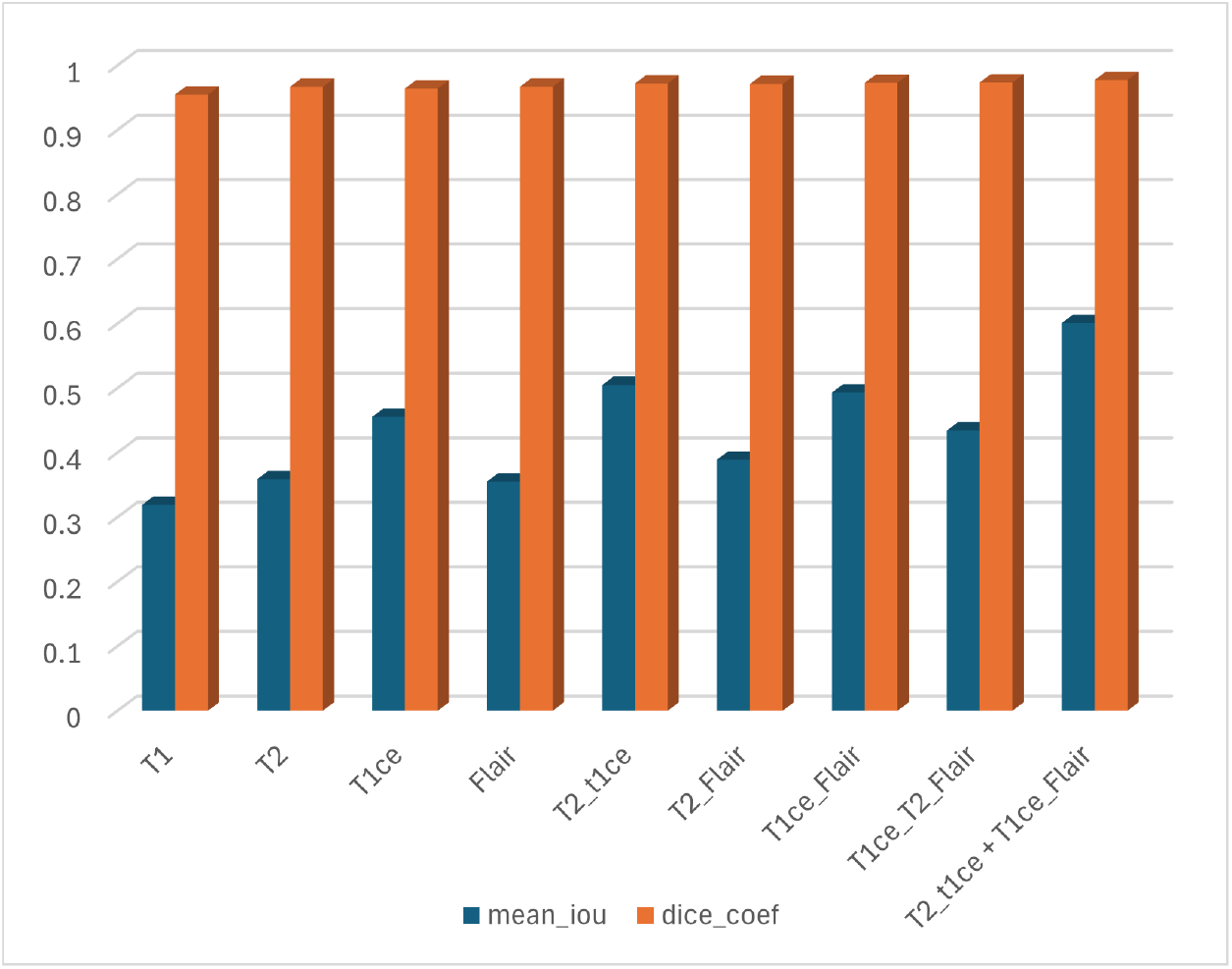
Results of Mean IoU and DSC for all scenarios.

### Single-Modality Segmentation

In this scenario, each MRI modality (T1, T2, T1ce, and FLAIR) was trained and tested on the U-Net model. The brain tumor segmentation performance was diverse among different MRI modalities during training and testing. The T1 modality achieved a Dice Coef of 0.954508 and a Mean IoU of 0.318932. Meanwhile, the T2 modality showed a Dice Coef of 0.966633 and a Mean IoU of 0.358804. In contrast, the T1ce modality obtained a Dice Coef of 0.964015 and a Mean IoU of 0.455298; T1ce images enhanced the visibility of the tumor due to contrast enhancement. Finally, the FLAIR modality resulted in a Dice Coef of 0.966846 and a Mean IoU of 0.355311. T1ce demonstrated the highest Mean IoU, indicating its potential for better tumor delineation. In addition, the T2 and FLAIR modalities performed better than the T1 modality. These results indicate that while single modalities provide valuable information, they have limitations in accurately capturing the full extent of the tumor.

### Multi-Modality Segmentation

U-Net models were trained using combinations of multiple MRI modalities. Combining modalities improved segmentation accuracy significantly, as shown in Table 1 and Figure 8. The T2 with T1ce modality achieved a Dice Coef of 0.971875 and a Mean IoU of 0.504634. While T2 with FLAIR, this combination resulted in a Dice Coef of 0.971087 and a Mean IoU of 0.388971. In contrast, the T1ce with FLAIR Has a Dice Coef of 0.972989 and a Mean IoU of 0.493028. Conversely, the T2 + T1ce + FLAIR triple-modality achieved a Dice Coef of 0.976191 and a Mean IoU of 0.579291. This multi-modality approach leveraged the complementary information provided by different MRI sequences, leading to more comprehensive tumor segmentation.

### Ensemble Dual-Modality Segmentation

The Ensemble Dual-Modality model combined the pre-trained models of the two best-performing dual-modality combinations (T1ce+FLAIR and T2+FLAIR). The Ensemble Dual-Modality model outperformed both the single-modality and multi-modality models by achieving a Dice Coef of 0.977302 and a Mean IoU of 0.600833. The Ensemble Dual-Modality model provided more accurate tumor segmentation because it integrated features from both dual-modality models, leveraging the strengths of each model. Also, using additional convolutional layers in the Ensemble Dual-Modality model helped merge features from different models, thus improving performance. Figure 9 shows an example of the result of the Ensemble Dual-Modality model applied.

**Figure 9.**
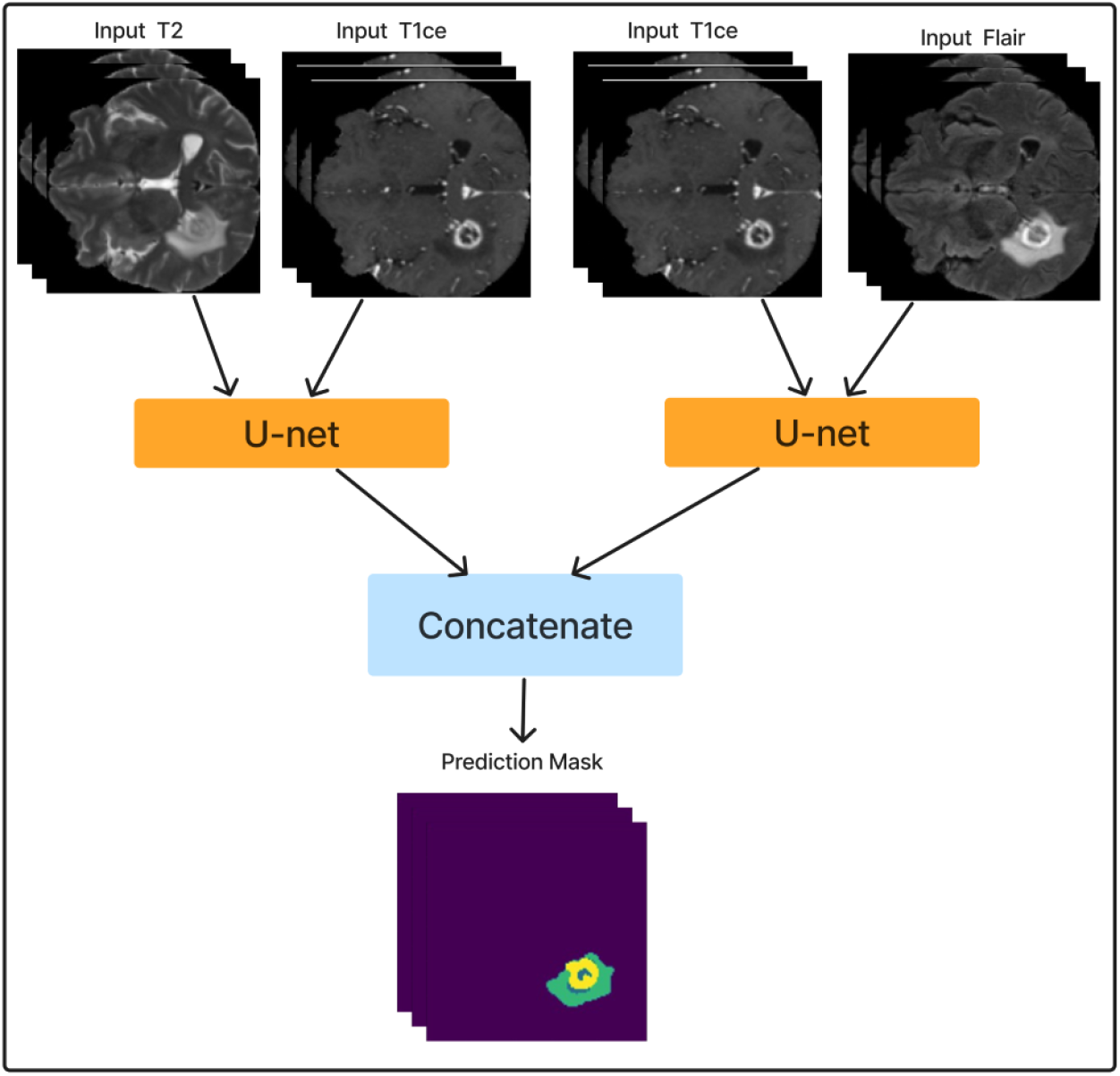
Example of applying the Ensemble Dual-Modality method

## DISCUSSION

The results highlight the advantages of using multi-modality MRI and an ensemble dual-modality approach for detecting and segmenting brain tumors. The results showed that using a single modality achieved different results because each modality highlights a specific aspect of the tumor characteristics. Therefore, combining more than one modality led to a significant improvement in the results. The combination of T2 + T1ce and T1ce + FLAIR methods outperformed the multi-modality and single-modality. This improvement is attributed to the complementary information of each modality and to the feature map of the model, which has become more complex.

The proposed ensemble dual-modality model, which combined two of the dual-modality models that obtained the best results, enhanced the model’s performance in segmenting brain tumours. The superior performance of the ensemble model, as demonstrated by the highest Dice coefficient and average IoU values, underscores its ability to leverage the strengths of each method to provide a more comprehensive and accurate tumor segmentation. In addition, the visualizations in Figures 10 to 18 in the Appendix illustrate the detailed segmentation outputs across various modalities and provide further evidence of our segmentation method’s robustness.

**Figure 10.**
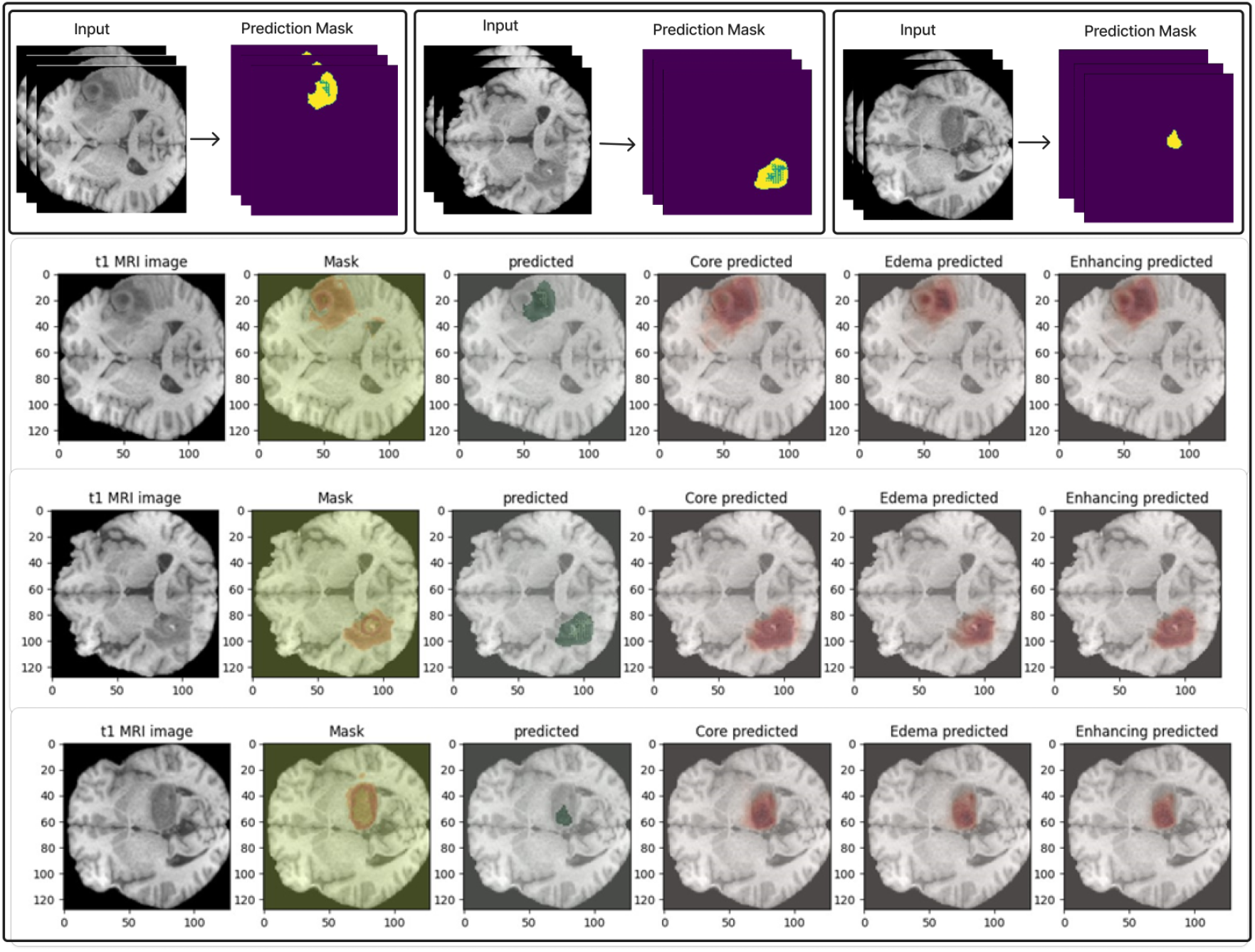
Example of segmentation results With just T1 modality input.

**Figure 11.**
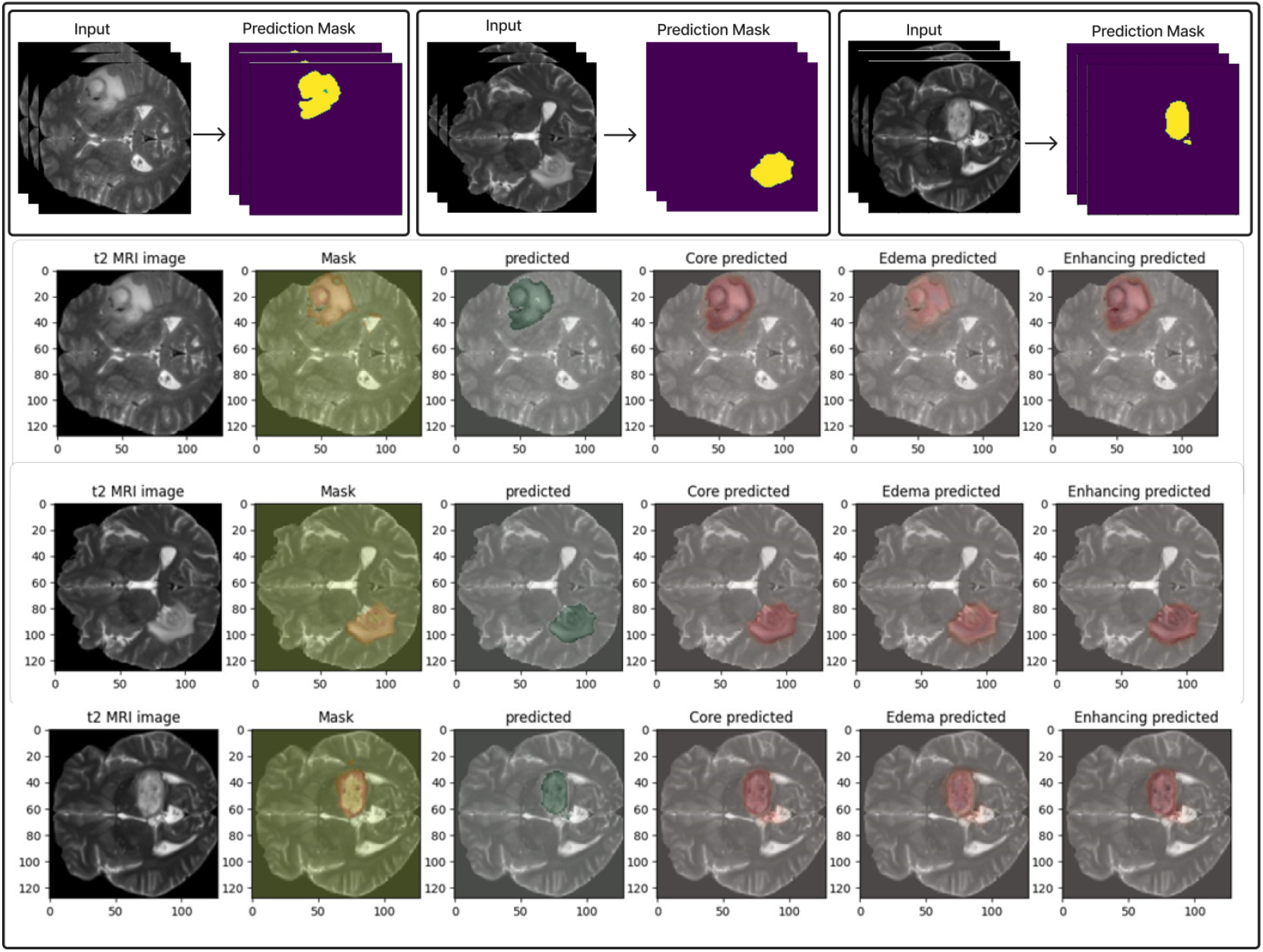
Example of segmentation results With just T2 modality input.

**Figure 12.**
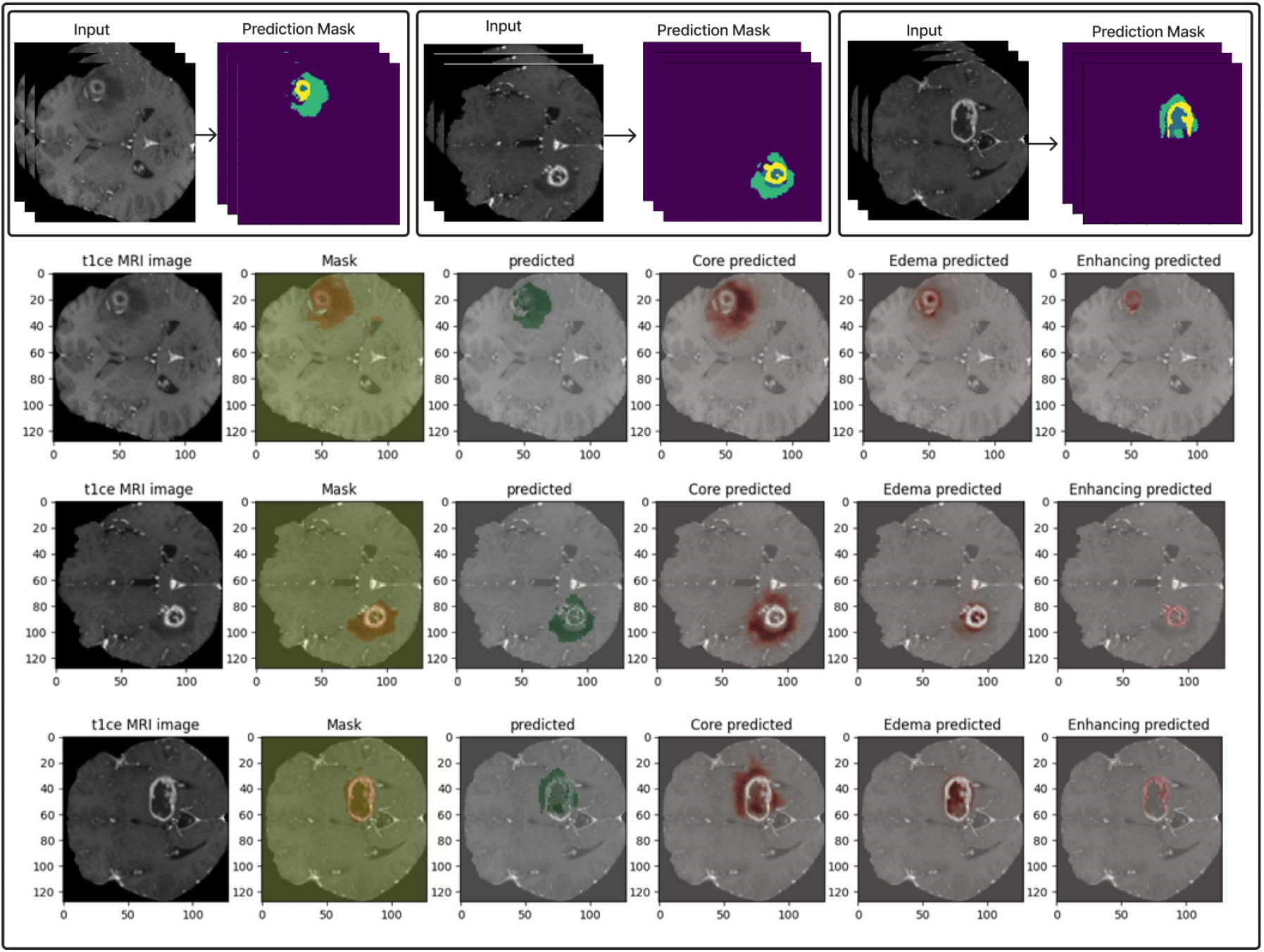
Example of segmentation results With just T1ce modality input.

**Figure 13.**
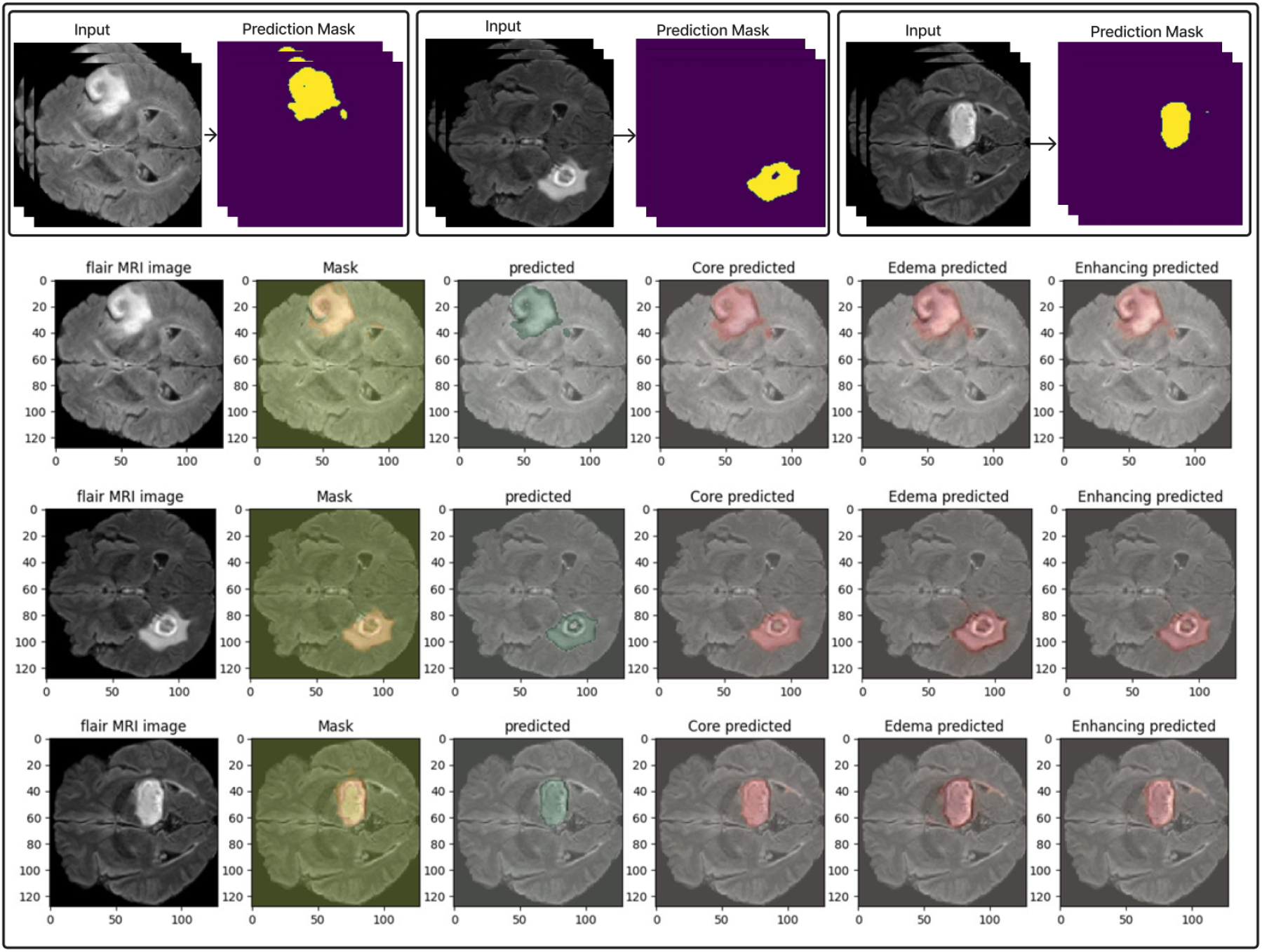
Example of segmentation results With just Flair modality input.

**Figure 14.**
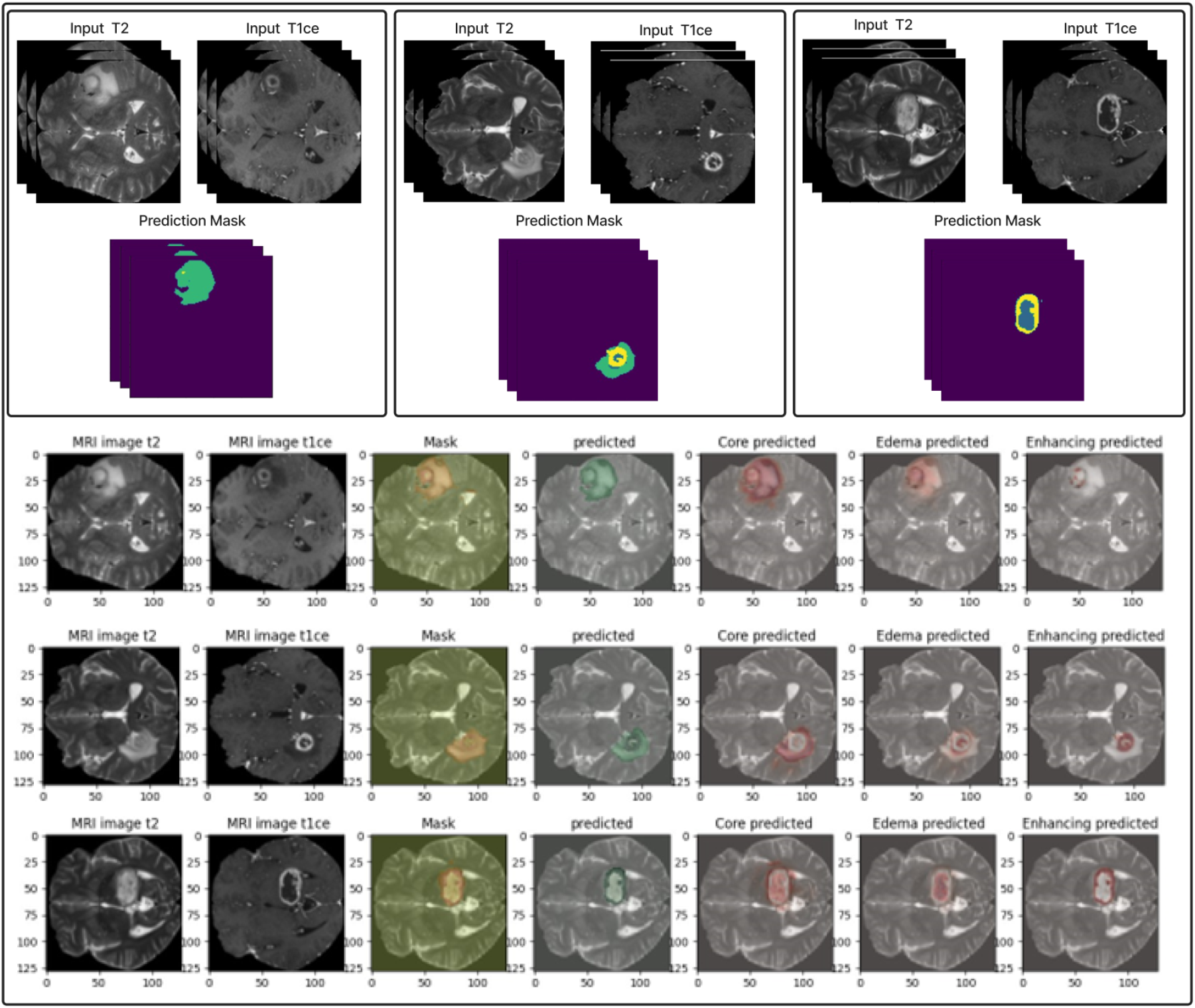
Example of segmentation results With T2 + T1ce modalities input.

**Figure 15.**
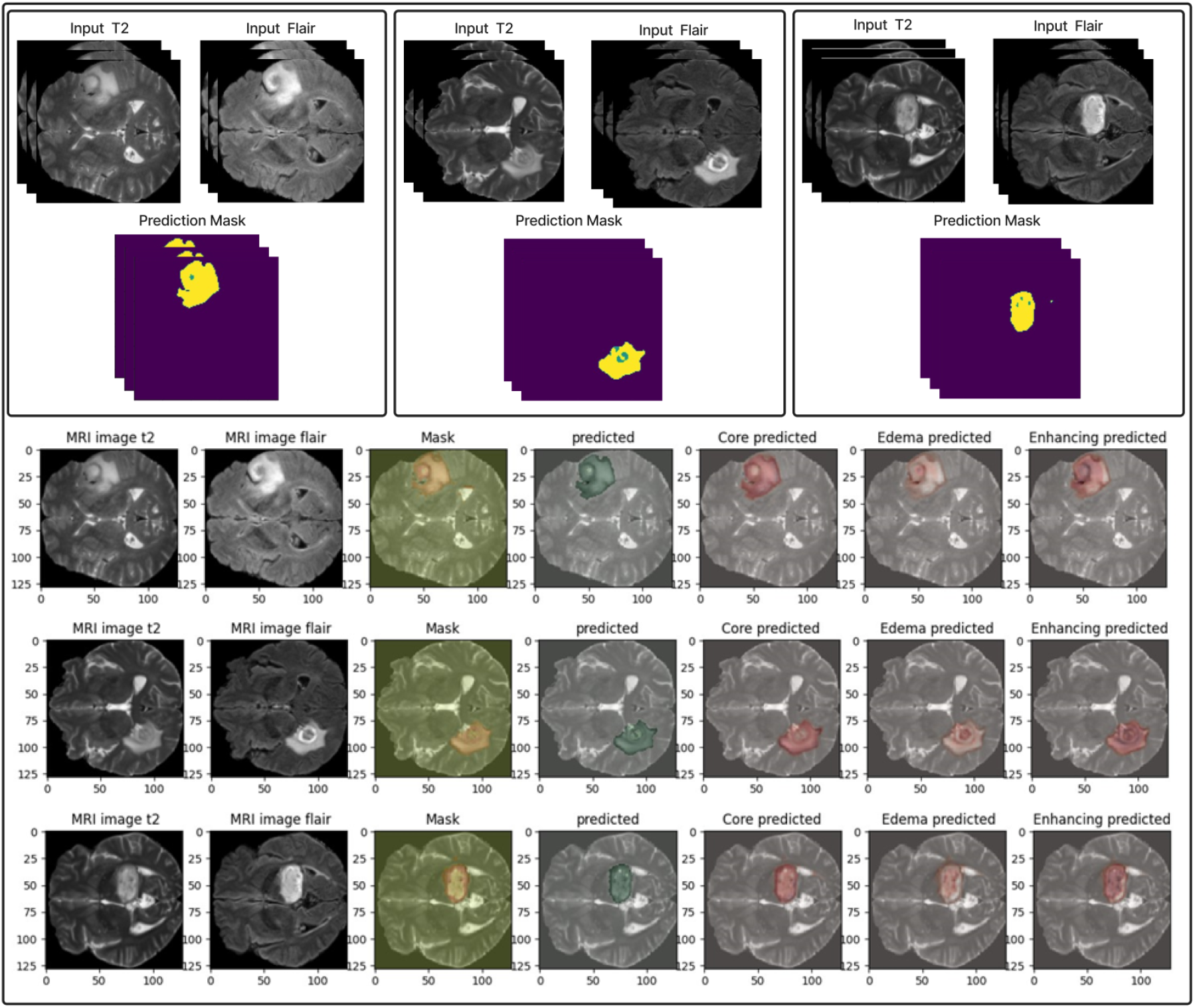
Example of segmentation results With T2 + Flair modalities input.

**Figure 16.**
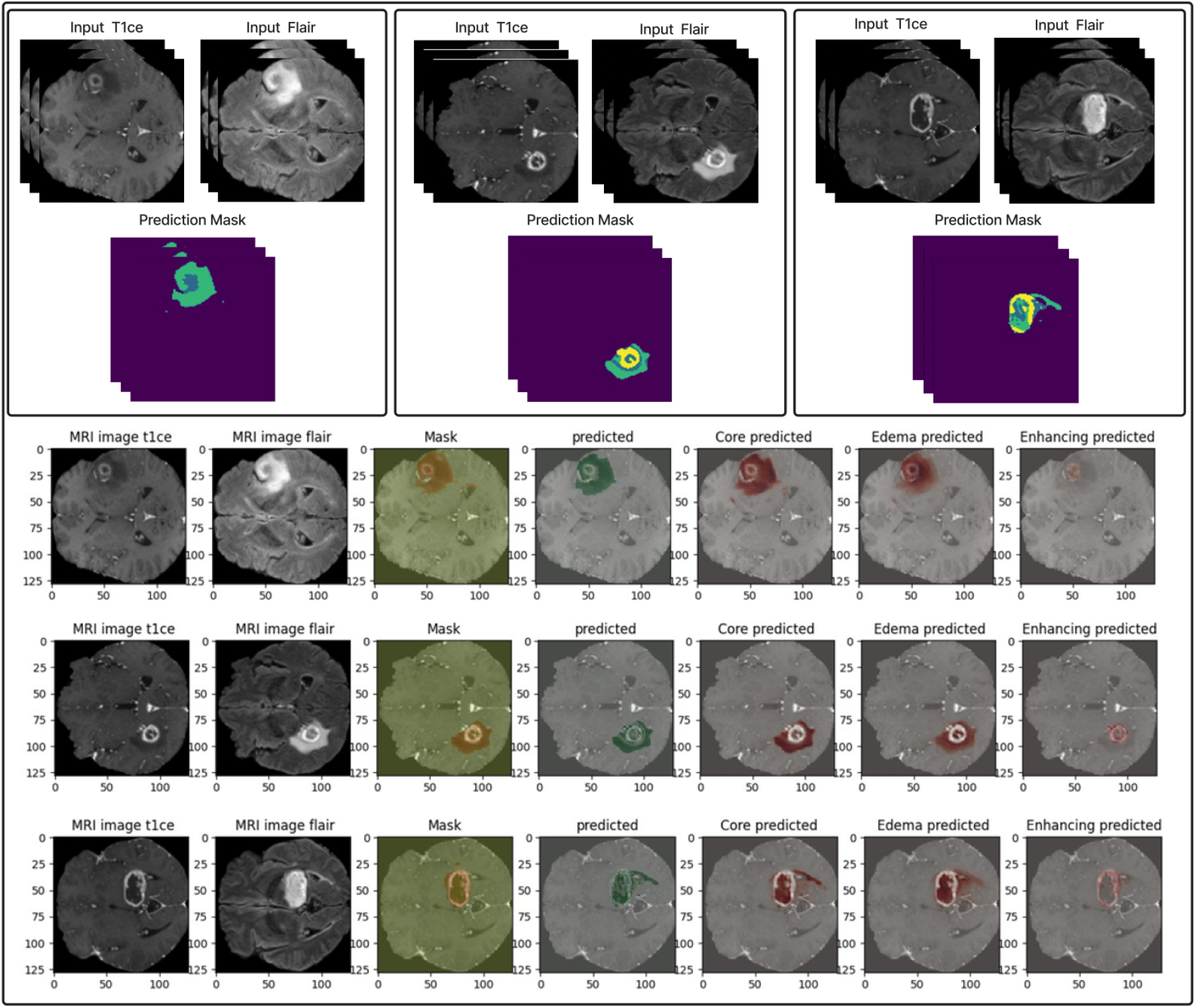
Example of segmentation results With T1ce+ Flair modalities input.

**Figure 17.**
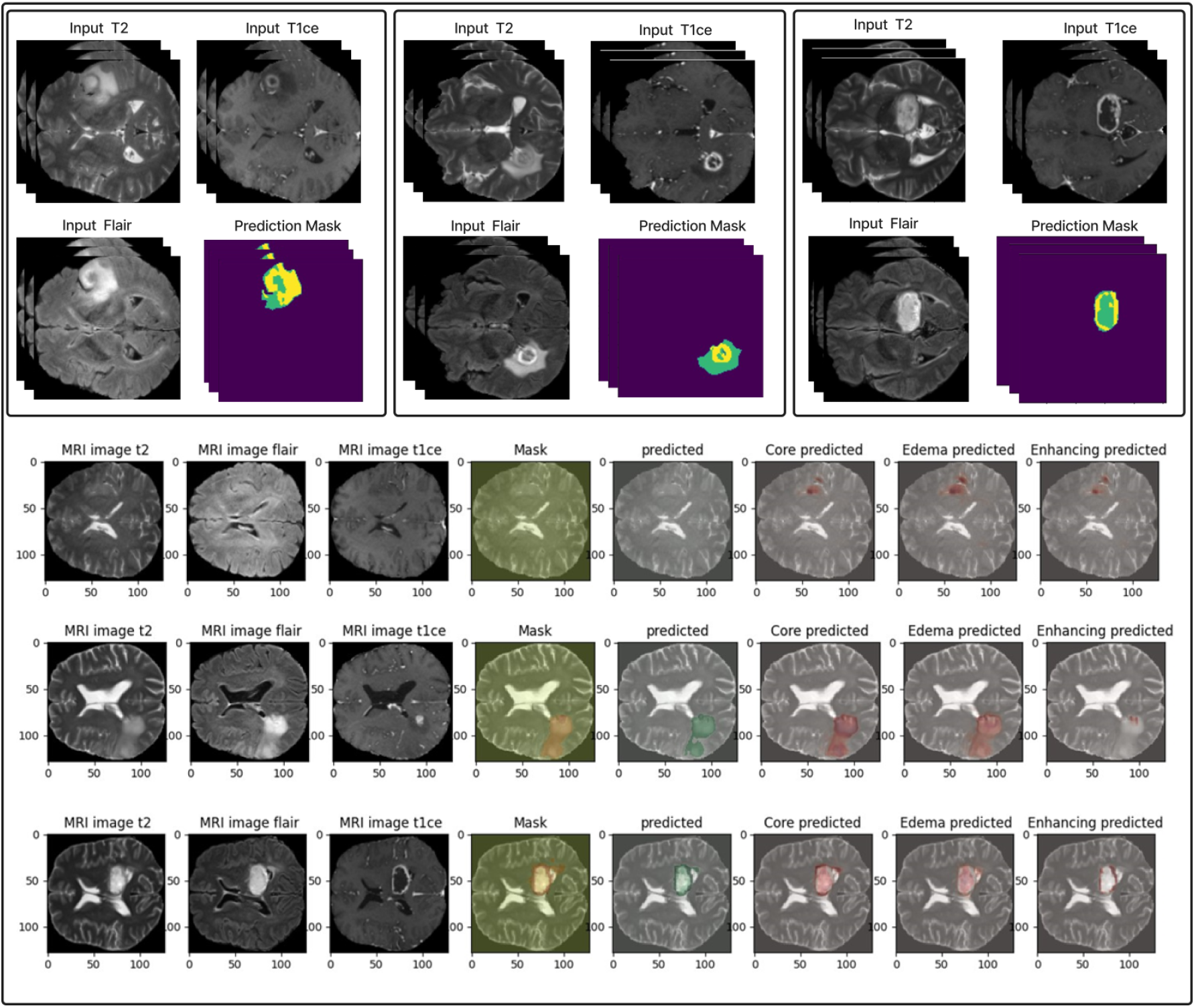
Example of segmentation results With T2 + T1ce + Flair modalities input.

**Figure 18.**
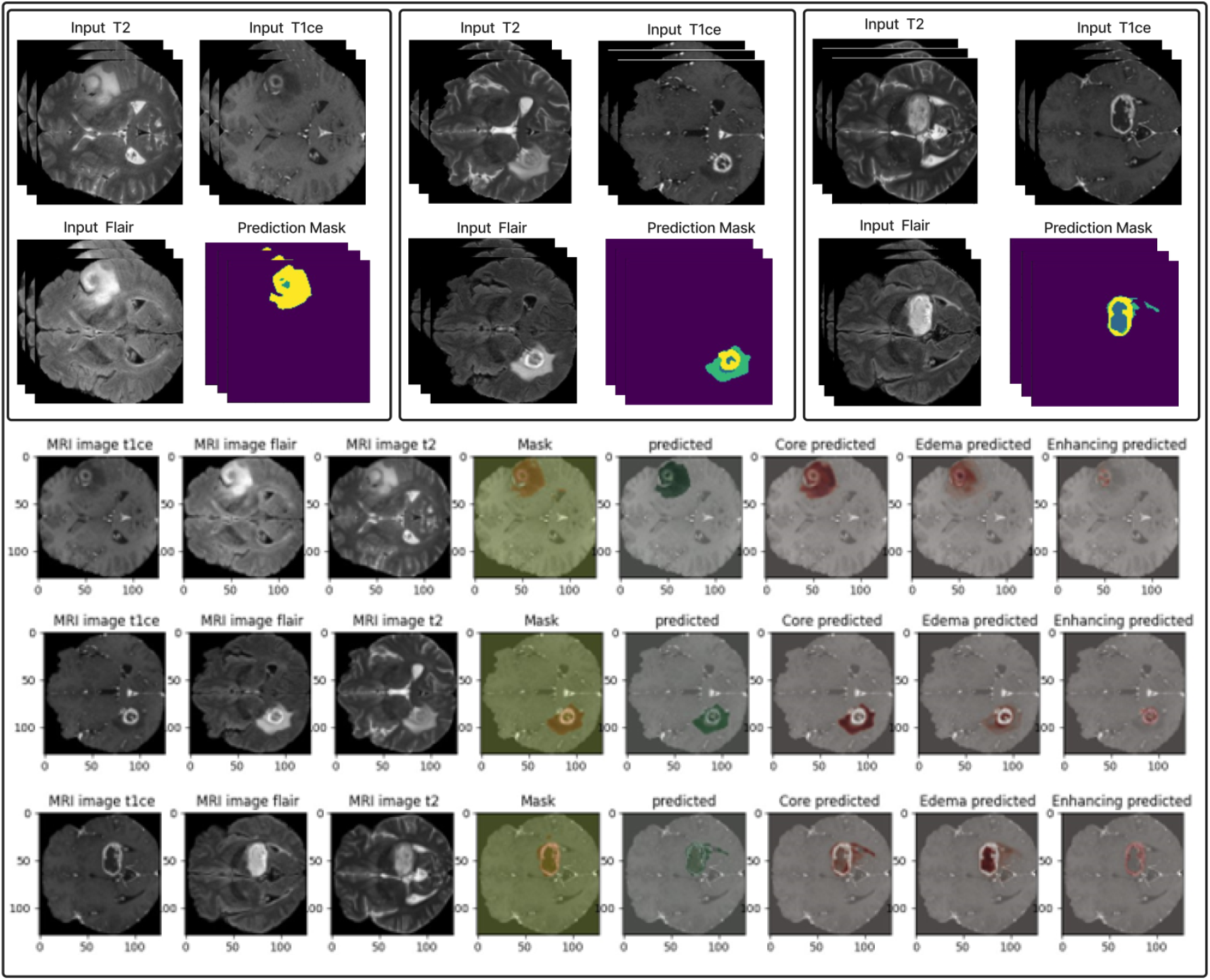
The Segmentation results example of Ensemble Dual-Modality approach

This study demonstrates the advantages of using multiple MRI modalities and ensemble approaches. However, there are limitations to consider. Integrating multiple modalities and ensemble models increases computational complexity and training time. Future work could focus on optimizing these models to reduce computational overhead while maintaining high segmentation accuracy. Additionally, further validation on larger and more diverse datasets could provide more generalized insights into the effectiveness of our proposed method.

## CONCLUSION

Brain tumor segmentation from Magnetic Resonance Images (MRI) presents significant challenges due to the complex nature of brain tumor tissues. This complexity makes distinguishing tumor tissues from healthy tissues difficult, mainly when radiologists perform manual segmentation. Reliable and accurate segmentation is crucial for effective tumor grading and subsequent treatment planning. Different MRI sequences such as T1, FLAIR, T1ce, and T2 provide unique insights into various aspects of the tumor. Our study proposed a novel ensemble dual-modality approach for 3D brain tumor segmentation from MRI. The proposed approach leverages the strengths of multiple MRI modalities and ensemble learning. The results demonstrate that while effective the U-net model with single-modality input can be significantly enhanced through dual-modality and ensemble methods. By combining T2, T1ce, and FLAIR modalities, dual-modality achieved better performance than single-modality in terms of Dice Coefficient and Mean IoU, underscoring the value of utilizing complementary information from different imaging techniques. The ensemble dual-modality model combined the two dual-modality pre-trained models that achieved the best results. The proposed approach achieved a Dice coefficient of 97.73% and a Mean IoU of 60.08% by testing on the BraTS2020 dataset. The proposed method leverages the strengths and characteristics of each modality to obtain accurate segmentation. The results also indicate the potential of leveraging ensemble learning for the purpose of segmenting medical images. Future work should improve the computational efficiency of the model, test it on a larger dataset, and validate the results by testing it in real-world applications in the clinic.

## APPENDICES

## Notes

### Competing Interest Statement

The authors have declared no competing interest.

